# Dorsal raphe stimulation relays a reward signal to the ventral tegmental area via GluN2C NMDA receptors

**DOI:** 10.1101/2020.07.26.222125

**Authors:** Giovanni Hernandez, Willemieke M Kouwenhoven, Poirier Emmanuelle, Lebied Karim, Daniel Lévesque, Pierre-Paul Rompré

## Abstract

**Background:** Glutamate relays a reward signal from the dorsal raphe (DR) to the ventral tegmental area (VTA). However, the role of the different subtypes of N-methyl-D-aspartate (NMDA) receptors is complex and not clearly understood. Therefore, we measured NMDA receptors subunits expression in limbic brain areas. In addition, we studied the effects of VTA down-regulation of GluN2C NMDA receptor on the reward signal that arises from DR electrical stimulation.

**Methods:** Using qPCR, we identified the relative composition of the different Grin2a-d subunits of the NMDA receptors in several brain areas. Then, we used fluorescent *in situ* hybridization (FISH) to evaluate the colocalization of Grin2c and tyrosine hydroxylase (TH) mRNA in VTA neurons. To assess the role of GluN2C in brain stimulation reward; we downregulated this receptor using small interfering RNA (siRNA) in rats self-stimulating for electrical pulses delivered to the DR. To delineate further the specific role of GluN2C in relaying the reward signal, we pharmacologically altered the function of VTA NMDA receptors by bilaterally microinjecting the NMDA receptor antagonist PPPA.

**Results:** We identified GluN2C as the most abundant subunit of the NMDA receptor expressed in the VTA. FISH revealed that about 50% of TH-positive neurons colocalize with Grin2c transcript. siRNA manipulation produced a selective down-regulation of the GluN2C protein subunit and a significant reduction in brain stimulation reward. Interestingly, PPPA enhanced brain stimulation reward, but only in rats that received the nonactive RNA sequence.

**Conclusion:** The present results suggest that VTA glutamate neurotransmission relays a reward signal initiated by DR stimulation by acting on GluN2C NMDA receptors.

**Highlights:**

- With the use of qpCR we identified the NMDA receptor composition in different brain areas
- Using Double-fluorescent in situ hybridization we demonstrated that TH+ cells contain the NMDA Glun2C subunit
- Using deep brain stimulation of the dorsal raphe and small interfering RNA we demonstrate that the reward signal is carried to VTA neurons through activation of Glun2C NMDA receptors.

## Introduction

The dorsal raphe nucleus (DR) plays a key role in the transmission of a reward signal [1–5]. It contains a heterogeneous group of neurons that send one of the highest projection density to the ventral tegmental area (VTA) [6], the origin of dopamine (DA) pathways implicated in reward and motivation [7–12]. Studies combining psychophysical methods with brain stimulation reward (BSR) revealed that activation of DR neurons triggers a strong reward signal [13,14] that travels along axons linking the DR to the VTA [3]; the combination of BSR with in vivo electrophysiology also showed that the DR reward signal activates a subpopulation of VTA DA-like neurons and that the magnitude of this activation is proportional to the magnitude of the reward signal [3,15]. The DR contains a large number of serotoninergic neurons [16], but it also contains glutamatergic, nitric oxide synthase, GABAergic and dopaminergic neurons [17]. The use of paired-pulse stimulation strongly suggests that serotonergic DR neurons do not carry the reward signal because the refractory periods of the reward-relevant neurons that linked DR to VTA are incompatible with those of serotoninergic neurons [3,18,19]. Indeed, more recent optogenetic studies revealed that the selective activation of DR glutamatergic and their VTA terminals triggers a reward signal [1,2,20]. Glutamate-containing terminals established asymmetrical synapses with DA neurons [2,21], and VTA glutamate release stimulates DA activity leading to the enhancement of extracellular DA in the nucleus accumbens, a limbic region that plays a key role in reward and motivation [2,22].

According to data from Qi et al. (2014), glutamate ionotropic AMPA receptors participate in the transmission of the DR reward signal to VTA mesoaccumbens DA neurons; however, several well-known characteristics of the interaction between glutamate and DA in the VTA suggest an important contribution of glutamate NMDA receptor subtypes (NMDAR). Activation of VTA NMDARs makes DA neurons to switch from a tonic, single-spike irregular firing pattern to a phasic burst firing pattern (Chergui et al., 1993a; Grace & Bunney, 1984; Hyland, Reynolds, Hay, Perk, & Miller, 2002; Johnson, Seutin, & North, 1992; P. G. Overton & Clark, 1997; Parker et al., 2010; Zweifel et al., 2009), a firing mode associated with an increased DA release and transmission of a reward signal [26,27]. But, besides this firing transition role relevant to reward, VTA glutamate also plays a role in maintaining a DA inhibitory drive (Grace et al., 2007). For example, pharmacological blockade of VTA NMDARs was shown to stimulate DA impulse flow and release, induce reward, and facilitate reward search [5]; an opposite effect is observed following the blockade of VTA NMDRs. The enhancement of DA neurotransmission and reward may occur via the attenuation of the tonic inhibitory afferent inputs to DA, increasing the probability that DA neurons fire in a burst mode. The opposite modulation of DA by glutamate may be explained by its action in different VTA neuronal populations expressing different subtypes of NMDAR. This hypothesis predicts that activation of a given subtype(s) potentiates DA burst firing and DA release, while activation of different subtype(s) increases the inhibitory drive to DA neurons. This idea is supported by data mentioned above and others showing that both activation and blockade of VTA NMDARs increase DA burst firing (French et al., 1993), accumbens DA release (Karreman et al., 1996; Westerink et al., 1996; Mathé et al., 1998; Kretschmer, 1999), and stimulate forward locomotion (Kretschmer, 1999; Cornish et al., 2001). Rodents also readily learn to directly self-administer the non-selective NMDAR antagonists, AP-7, into the VTA (David et al., 1998), showing that NMDAR blockade can have positive, rewarding properties on its own.

NMDARs are formed by tetrameric complexes composed of two obligatory GluN1 subunits that cluster with GluN2 and/or GluN3 subunits [28,29]. In a previous study of our group, we reported that down-regulation of GluN1 subunit protein in VTA neurons, which produces a non-selective reduction in NMDAR expression, results in a specific attenuation of reward induced by DR electrical stimulation [30]. However, we could not identify the GluN1 subunit partner because a similar downregulation of the GluN2A or GluN2D subunit failed to alter brain stimulation reward [30]; Moreover, selective pharmacological blockade of VTA GluN2B subunits did not affect the reward signal arising from the DR [5].

Here, we first report that Grin2c mRNA is expressed in the VTA in higher concentration than other Grin2 transcripts (a, b, or d) and colocalizes within tyrosine hydroxylase positive (TH+) and negative (TH-) neurons, and second that selective downregulation of VTA NMDAR containing GluN2C subunits leads to a specific attenuation of the reward signal initiated by DR electrical stimulation.

## Materials and Methods

### Subjects and surgery

Twenty-two (22) male Long-Evans rats (Charles River, St-Constant, QC) weighing between 350-400 g at the time of surgery were used. Rats were individually housed in a temperature- and humidity-controlled room with a 12-h light-dark cycle (lights on at 06:00 h) and *ad libitum* access to food and water. After a minimum 7-day period of acclimatization to the housing, environment rats were anesthetized with isoflurane (2.5-3.5% O2, 0.6 L/min) and stereotaxically implanted according to the coordinates of Paxinos and Watson (2007) with 26-gauge guided cannulae (HRS Scientific, Montreal, Canada) aimed bilaterally at the VTA (−5.5 AP, 3.2 ML at an angle of 18 °, 6.5 DV from the skull surface) and a monopolar stainless steel monopolar electrode aimed at the DR (−7.6 AP, 0 ML, 6.6 DV from the skull surface). Detailed surgical procedures can be found in Bergeron and Rompré (2013). All procedures were approved by the Animal Care and Use Committee of the Université de Montréal in accordance with the guidelines of the Canadian Council on Animal Care.

### siRNAs and Drugs

Downregulation of selective NMDAR subunit expression was achieved using a pre-validated small interfering RNA (siRNA) sequence against the rat glutamate ionotropic receptor NMDA type subunit 2C (Grin2c) mRNA, which encodes the GluN2C protein subunit (Silencer™ select pre-designed siRNA # 4390771, siRNA ID s127815, ThermoFisher Scientific) and a nonactive RNA sequence (Silencer™ select negative control #4390844, ThermoFisher Scientific). The deprotected, duplexed, desalted siRNA was mixed with a cationic lipid transfection carrier N-[1-(2,3-Dioleoyloxy)propyl]-N, N, N-trimethyl-ammonium methylsulfate (DOTAP) (Roche Applied Sciences, Indianapolis, IN), which showed high efficacy for *in vivo* transfection [31]. The final solution contained 10 μg of active or inactive siRNA and 1 μg of DOTAP per μl.

PPPA ((2R, 4S) -4- (3-phosphatophores) -2-piperidinecarboxylic acid), a competitive GluN2A-favored NMDAR antagonist (Tocris, Ellisville, MI, USA), was dissolved in 0.9% sterile saline and stored frozen in 40–50 μl aliquots. Drug solutions were thawed just before testing and used only once. PPPA was injected into VTA at a dose of 0.825 nmol/0.5μl/side, as previously described [5,30,32].

### Self-stimulation training

Each rat was trained to nose-poke for a 0.4-sec train of cathodal, rectangular, constant-current pulses, 0.1 msec in duration, delivered at a frequency of 98 Hz. For a detailed shaping procedure, see [5]. Once the rat nose-poked consistently for currents between 125 and 400 μA, a rate vs. pulse-frequency curve was obtained by varying the stimulation frequency across trials over a range that drove the number of rewards earned from maximal to minimal levels. The stimulation frequency was decreased from trial to trial by approximately 0.05 log10. Each trial for obtaining the rate-frequency sweep lasted for 55 s, followed by a 15 s inter-trial interval during which stimulation was not available. The beginning of each trial was signaled by five trains of non-contingent priming stimulation delivered at a rate of 1 per second. Four sweeps were run daily, and the first sweep was considered a warm-up and discarded from the analysis. The data relating to the rate-frequency was fitted to a sigmoid described by the following equation y=Min+((Max-Min))/(1+[10]^((x50-x)*p)) where Min is the lower asymptote, Max is the upper asymptote, x50 is the position parameter denoting the frequency at which the slope of the curve is maximal, and p determines the steepness of the sigmoid curve. The resulting fit was used to derive an index of reward defined as the pulse-frequency sustaining a half-maximal rate of responding (M50). Self-stimulation behavior was considered stable when the M50 values varied less than 0.1 log unit for three consecutive days.

Prior to the siRNA injections or drug administration, sterile 0.9% saline was microinjected into the VTA to habituate the animals to the injection procedure. Bilateral injections were performed by inserting an injection cannula (Model C315I HRS Scientific, Montreal, Canada) that extended 2 mm beyond the tip of the guide cannula. The injection cannula was connected to a 5 μl Hamilton microsyringe via polyethylene tubing. A total volume of 0.5 μl was injected into each hemisphere simultaneously over a 60-s period. The rate of delivery was controlled by an infusion pump (Harvard Instruments, Holliston, MA). The injection cannulae were left in place for an additional 60 s to allow diffusion into the tissue. After microinjection, the subjects were put into the operant boxes and allowed to self-stimulate. The results of this test were not included in the analysis. Baseline data were collected one week after this first saline microinjection. Once a stable baseline was obtained, pre-validated siRNA (5 μg per side) against GluN2C or nonactive RNA sequence was injected bilaterally into VTA for two consecutive days. Reward thresholds were measured 24-h after each injection. After 24 h, the last threshold determination rats received a bilateral VTA microinjection of the NMDA antagonist, PPPA, and reward thresholds were measured again immediately after injection for 120 min (see supplementary Fig. S1 for experiment timeline).

### Western Blot

Rats were decapitated immediately after the last behavioral test. Brains were removed and directly placed on an ice-cold brain matrix and sectioned coronally. The VTA was dissected on an ice-cooled plate from a 0.75-1mm slice using a 15-gauge tissue punch. The tissue was placed in a 1 ml Eppendorf container and then immediately frozen at -80°C until biochemical experiments were performed. VTA samples were mechanically homogenized in lysis buffer [10 mM Tris–HCl (pH 6.8), 2% SDS and a cocktail protease inhibitor (Roche)]. The protein concentrations of the tissue samples were measured using the BCA™ protein assay kit (Pierce, USA). Equal amounts of protein (10 μg) were dissolved into 25 μl lysis buffer (which contained 4.5 μl 5X loading buffer and 0.5 μl β-mercaptoethanol), boiled at 95°C for 5 min, and then subjected to SDS polyacrylamide gel electrophoresis (PAGE) with 8% polyacrylamide and transferred to polyvinylidene difluoride (PVDF) membrane (BioRad Laboratories). The membranes were blocked for 1 h in Tris buffered saline with Tween 20 (TBST) buffer with 5% bovine serum albumin (BSA) and incubated with primary rabbit anti-β-actin antibody (Millipore Sigma, cat. #SAB5600204) at 1:20000, with either primary mouse anti-GluN2C antibody (Novus Biological, cat. no. #NBP2-29809) or mouse anti-GluN2A antibody (Millipore, cat. #SAB5200897) at 1:500 overnight at 4°C. After rinsing 4 times with TBST for 5 min, the membranes were incubated with Horse Raddish Peroxidase (HRP)-conjugated goat anti-mouse IgG secondary antibody (Cell Signaling, cat. #7076) at 1:20000 (to detect GluN2C or GluN2A) or HRP-conjugated goat anti-rabbit IgG secondary antibody (Cell Signaling, cat. #7074) at 1:40000 (to detect β-actin) for 1 h. Membrane protein bands were detected with ECL (BioRad) and visualized on the Chemidoc Biorad system (BioRad). Band densities were measured and analyzed with Image Lab software (BioRad), and protein levels were normalized over β-actin. The different subunits of the NMDAR were expressed as a percentage of control.

### Double-fluorescent in situ hybridization (FISH)

Three naive Long-Evans male rats were anesthetized using an overdose of intraperitoneal ketamine 50⍰mg/kg, xylazine 5 mg/kg, and acepromazine 1⍰mg/kg perfused intracardially with ice-cold PBS. After removal, the brains were rapidly frozen in dry ice-cooled 2-methyl butane (Fisher Scientific, Hampton, NH, USA). Coronal sections of the VTA at 10⍰μm were obtained using a cryostat and mounted onto superfrost slides (Fisher Scientific Ltd, Nepean, ON, Canada). Coronal cryosections were prepared, air dried, fixed in 4% paraformaldehyde, and acetylated in 0.25% acetic anhydride/100 mM triethanolamine (pH 8), followed by incubation for 30 minutes at room temperature in hybe solution. Then, slices were incubated overnight at 60°C in 100 μl of hybe-solution buffer containing 20 ng digoxigenin (Dig)-labeled probes (see below) for colorimetric detection (Roche, cat. #11277073910) or 20 μg of fluorescein-labeled probes (see below) for fluorescent detection (Roche, cat. #11685619910). After hybridization, sections were washed at 60°C with SSC buffers of decreasing strength (5X-0.2X), then washed at room temperature with 0.2X SSC and maleic acid buffer containing tween 20 (MABT). After washing, sections were blocked for 30 minutes with 20% FBS and 1% blocking solution. For colorimetric detection, Dig epitopes were detected with alkaline phosphatase-coupled anti-Dig fab fragments antibody at 1:2500 (Millipore Sigma, cat. #11093274910), and the signal revealed with Nitro Blue Tetrazolium chloride/5-Bromo-4-Chloro-3-Indolyl Phosphate, toluidine salt (NBT/BCIP) alkaline phosphatase chromogen substrate solution (SigmaAldrich).

For fluorescent detection, sections were incubated with HRP-conjugated anti-fluorescein antibody at 1:500 concentration. Signals were revealed with the Tyramide Signal Amplification (TSA plus biotin) kit (PerkinElmer, cat. #NEL749A001KT) at 1:100 concentration, followed by incubation with neutravidin Oregon Green conjugated at 1:500 (Invitrogen, cat. #A-6374). HRP activity was stopped by the incubation of sections in 0.1 M glycine and 3% H_2_O_2_. Dig epitopes were detected with HRP-conjugated anti-Dig antibody at 1:1000 (Roche cat. #11207733910) and revealed with the TSA kit (PerkinElmer, cat. #NEL704A001KT) using Cy3 tyramide at 1:100. All sections were counterstained with DAPI and mounted using Fluoromount (ThermoFisher Scientific, cat. #00-4958-02).

Double FISH was performed with RNA probes labeled with digoxigenin (Dig) for tyrosine hydroxylase and RNA probes labeled with Fluorescein for Grin2C, consisting of a 678 bp fragment, (Fw primer: 5’-ctactgctcccgtgaagagg-3’, Rv primer: 5’-agagcttggtgtagggggtt-3’). The tyrosine hydroxylase probe consisting of a fragment of 1142 bp was a kind gift from Prof. Marten Smidt (University of Amsterdam). Cells were automatically counted using ImageJ FIJI software (NIH, USA). Briefly, each channel was converted to binary and combined. This creates an image of only the cells that expressed both TH and NR2C. Then we automatically counted the numbers of cells in the TH binary and the number of cells in the TH+NR2C binary. The percentage was calculated by dividing the total number of TH+ cells expressing NR2C by the total number of TH+ cells and multiplying the number by 100.

### qRT-PCR analysis

Because no study has been carried out to determine the extent of the different NMDAR subunits in the limbic brain regions involved in reward and motivation, we carried out the following quantitative real-time RT-PCR study of the Grin2a-d transcript subtype. Brain punches from the prefrontal cortex (PFC), nucleus accumbens (Nac), lateral hypothalamus (LH), habenula (Hab), amygdala (Amyg), bed nucleus of the stria terminalis (BNst), ventral pallidum (VP), hippocampus (Hipp), pedunculopontine tegmental nucleus (PPtg), ventral tegmental area (VTA), rostromedial tegmental nucleus (RMtg) and raphe nucleus of four (n=4) naive Long-Evans male rats were taken. Total RNA was extracted using trizol and purified on mini spin columns (RNeasy Kit protocol, Qiagen, Toronto, ON, Canada). All RNA samples were determined to have values 260/280 and 260/230 values >1.8, using the Nanodrop 1000 system (Thermo Scientific, Toronto, ON, Canada). We assessed the integrity of RNA by the presence of well-defined 28S and 18S ribosomal bands and RNA integrity number ≥ 8 using the 2100 bioanalyzer (Agilent). Reverse transcription (RT) for Grin2a, Grin2b, Grin2c, Grin2d, glyceraldehyde-3-phosphatedehydrogenase (GAPDH), and hypoxanthine phosphoribosyltransferase (HPRT) was performed (see primer sequences in supplementary Table 1). Real-time PCR, using a TaqMan assay, was performed with an Applied Biosystems 7900HT RT PCR system on the Institute for Research on Immunology and Cancer (IRIC) genomic platform (https://genomique.iric.ca/). Data were analyzed using the relative quantification method (ΔΔ ct), and the levels of transcripts were normalized with both the expression of the reference (housekeeping) genes, GAPDH and HPRT. We used two housekeeping genes to make a robust determination of the levels of Grin2c across the brain areas studied. As expressed by Vandesompele et al.(2002), the use of multiple housekeeping genes significantly decreases the estimation error in qPCRs. When using multiple housekeeping genes for accurate normalization of the relative gene expression, the geometric mean is used in the calculations.

### Data Analysis

The fit of the rate frequency data and M50 value were obtained using MATLAB (Natick, MA). Differences in M50 values and protein levels were assessed using the t-test. Data analysis was performed using Statistica v12 (Tulsa, OK), and graphics were done in Origin v9 (Northamptom, MA).

## Results

### Expression of NMDAR subunit mRNAs (Grin2a-d) in the VTA and other limbic areas

Considering the role that glutamate and NMDAR play in modulating the limbic circuitry implicated in motivation and, particularly, in relaying a reward signal in the VTA, we first investigated the distribution of different NMDAR subunits throughout several limbic regions. We performed qRT-PCR on mRNA taken from all relevant regions (Fig. 1: the prefrontal cortex (PFC), nucleus accumbens (Nac), lateral hypothalamus (LH), habenula (Hab), amygdala (Amyg), bed nucleus of the stria terminalis (BNST), ventral pallidum (VP), hippocampus (Hipp), pedunculopontine tegmental nucleus (PPtg), ventral tegmental area (VTA), rostromedial tegmental nucleus (RMtg) and dorsal raphe nucleus (DR). As expected, mRNA concentrations vary notably across brain areas, although in all areas examined, we observed that the expression of the Grin2d subunit remained relatively low compared to other subunits. In Hipp, the mRNA for the Grin2a and Grin2b subunits are the most abundant, followed by Grin2c. Grin2b mRNA is the most abundant in the PFC, followed by Grin2a and Grin2c. Grin2b and Grin2c were the most abundant in the nucleus accumbens, the ventral pallidum and the BNst. In Amyg Grin2b mRNA was also the most abundant, but Grin2a was more elevated than Grin2c. A nearly similar content of Grin2b and 2c was observed in LH and Hab. Grin2c was the most abundant in regions posterior to the VTA, namely the PPTg, RMTg, and Raphe nuclei. We found that Grin2c mRNA is expressed within the VTA, being as abundant as 2b. We also confirmed that the Grin2c transcript is translated into a protein subunit (GluN2C) in the VTA (data not shown) by Western blotting.

**Figure 1.**
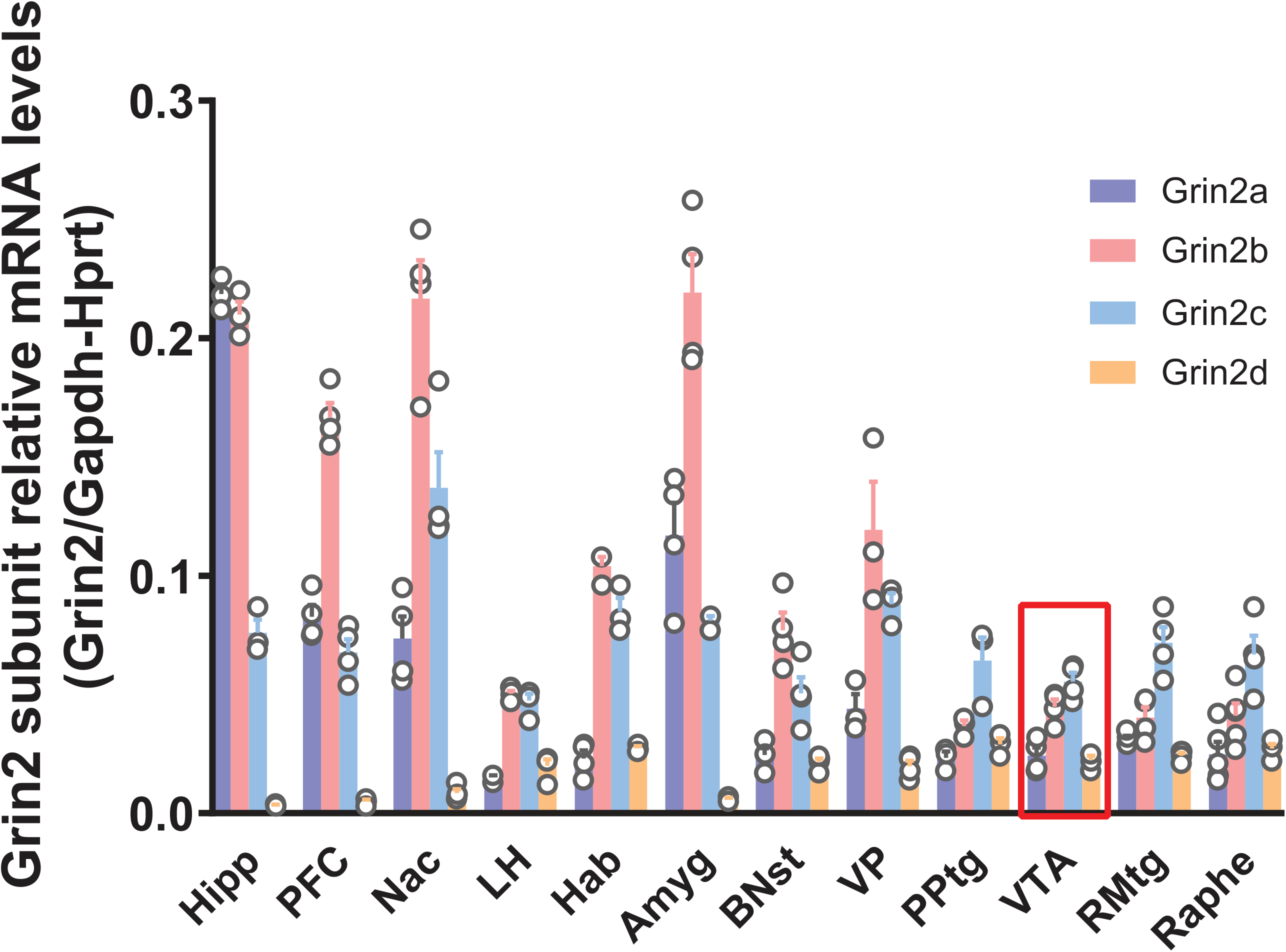
Relative expression of NMDAR containing Grin2a-d subunits mRNA across different brain areas. Brain punches from the prefrontal cortex (PFC), nucleus accumbens (Nac), lateral hypothalamus (LH), habenula (Hab), amygdala (Amyg), bed nucleus of the stria terminalis (BNST), ventral pallidum (VP), hippocampus (Hipp), pedunculopontine tegmental nucleus (PPtg), ventral tegmental area (VTA), rostromedial tegmental nucleus (RMtg) and raphe nucleus (Raphe were analyzed) (n=4). Relative mRNA levels were quantified using ΔΔ ct method. Histogram bars represent relative mRNA levels normalized over both GAPDH and HPRT mRNA levels.

To confirm and elaborate on the observation that VTA neurons express significant levels of Grin2c, we performed an independent set of experiments to investigate the spatial expression of Grin2c in the mesencephalon. The Allen Brain Atlas displays Grin2c expression in the mesencephalon, with expression in the red nucleus (arrowhead, Fig. 2A) and SN and VTA region (arrow, Fig 2A) [34,35]. Using the same probe, we confirmed the ample expression of Grin2c in the midbrain (Fig 2A). To verify if Grin2c is expressed in DA neurons, we performed double-labeling fluorescent *in situ* hybridization and observed co-expression of Th transcript and Grin2c in the VTA (Fig 2B). Quantitative analysis shows that the level of colocalization of Grin2c and TH in DA neurons is around 46% (SEM= 7.83) (Fig. 2C and supplementary Fig. S2). Together, these data suggest that Grin2c could be a relevant component in transmitting glutamatergic signals by DA VTA neurons.

**Figure 2.**
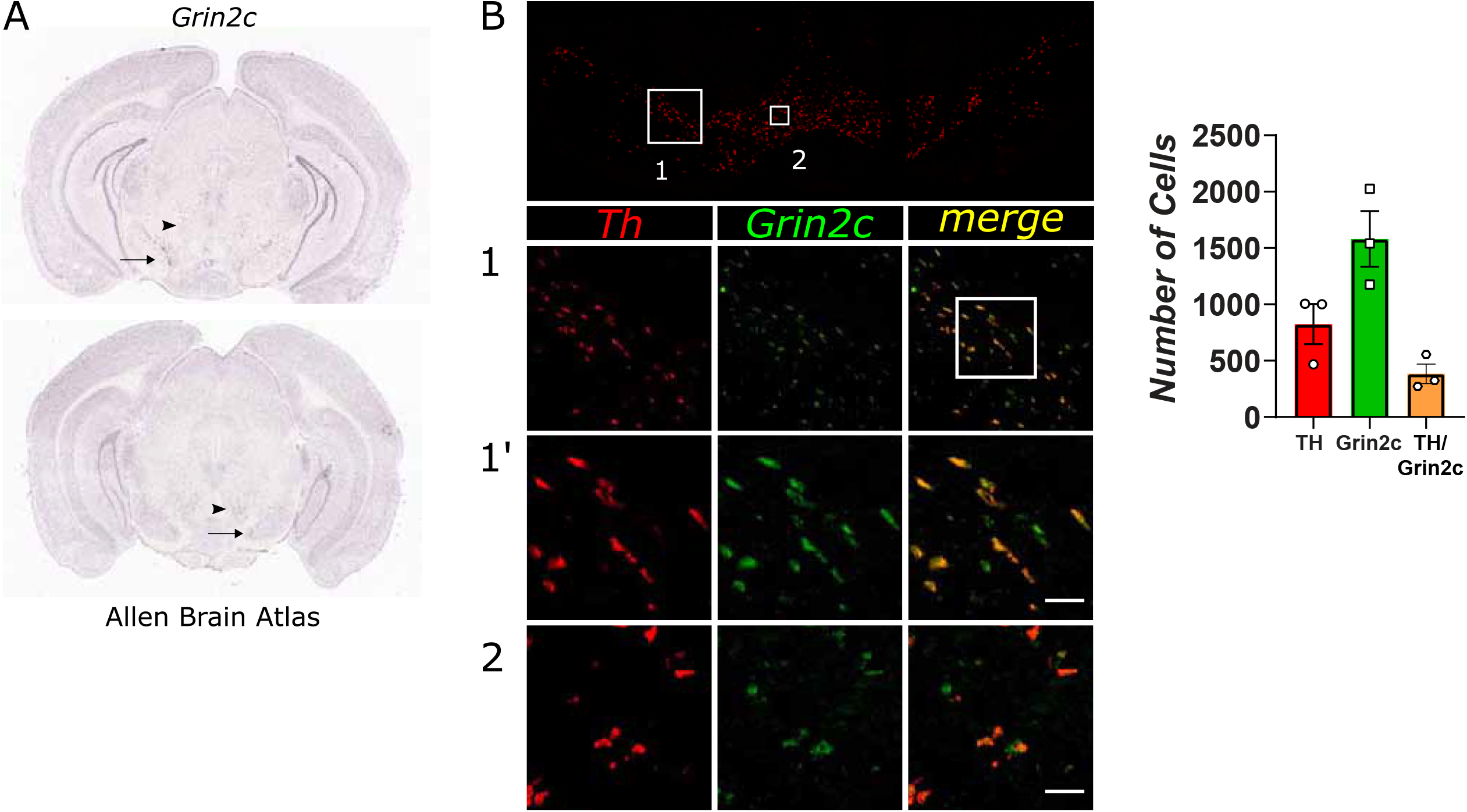
Double FISH immunodetection shows a high colocalization of Grin2c in TH positive cells of the VTA. **A**. Representative images of the ventral midbrain area containing the SN and VTA taken from the Allen Brain Atlas, displaying Grin2c transcript in the red nucleus (arrowhead) and SN and VTA area (arrow). B-B”. Fluorescent *in situ* hybridization reveals that Grin2c expression (Fluorescein; green) overlaps with tyrosine hydroxylase (TH) (Cy3; red) in the mesencephalon, scale represents 50 μm. **C**. Bar graphs showing the count of TH and Grin2c positive cells in the VTA.

### Validation of siRNA effects on NMDA receptor subunit protein levels

Knowing that the Grin2c is significantly expressed in VTA DA neurons, we set out to determine its involvement in relaying the glutamate reward signal. To do so, we designed a strategy to specifically down-regulate the NMDA receptor GluN2C protein subunit *in vivo* by injecting adult rats bilaterally with siRNA against Grin2c transcript, or scrambled siRNA as a control, in the VTA (−5.5 AP, 3.2 ML at an angle of 18 °, 6.5 DV from the skull surface). To confirm the efficacy and the specificity of the Grin2c siRNA, we performed Western blots for GluN2C and GluN2A in total homogenates of VTA (immediately after the last behavioral test). Microinjections of siRNA against Grin2c produced a significant 41% (SEM = 3.22) decrease in the expression of the NMDAR receptor GluN2C subunit compared to the scrambled control [t_(10)_ =2.43; p=0.035] (Fig. 3A). Moreover, the active Grin2c siRNA was selective for GluN2C subunit since we observed no change in protein expression of the GluN2A subunit [t_(10)_ =0.28; p=0.783] (Fig. 3B).

**Figure 3.**
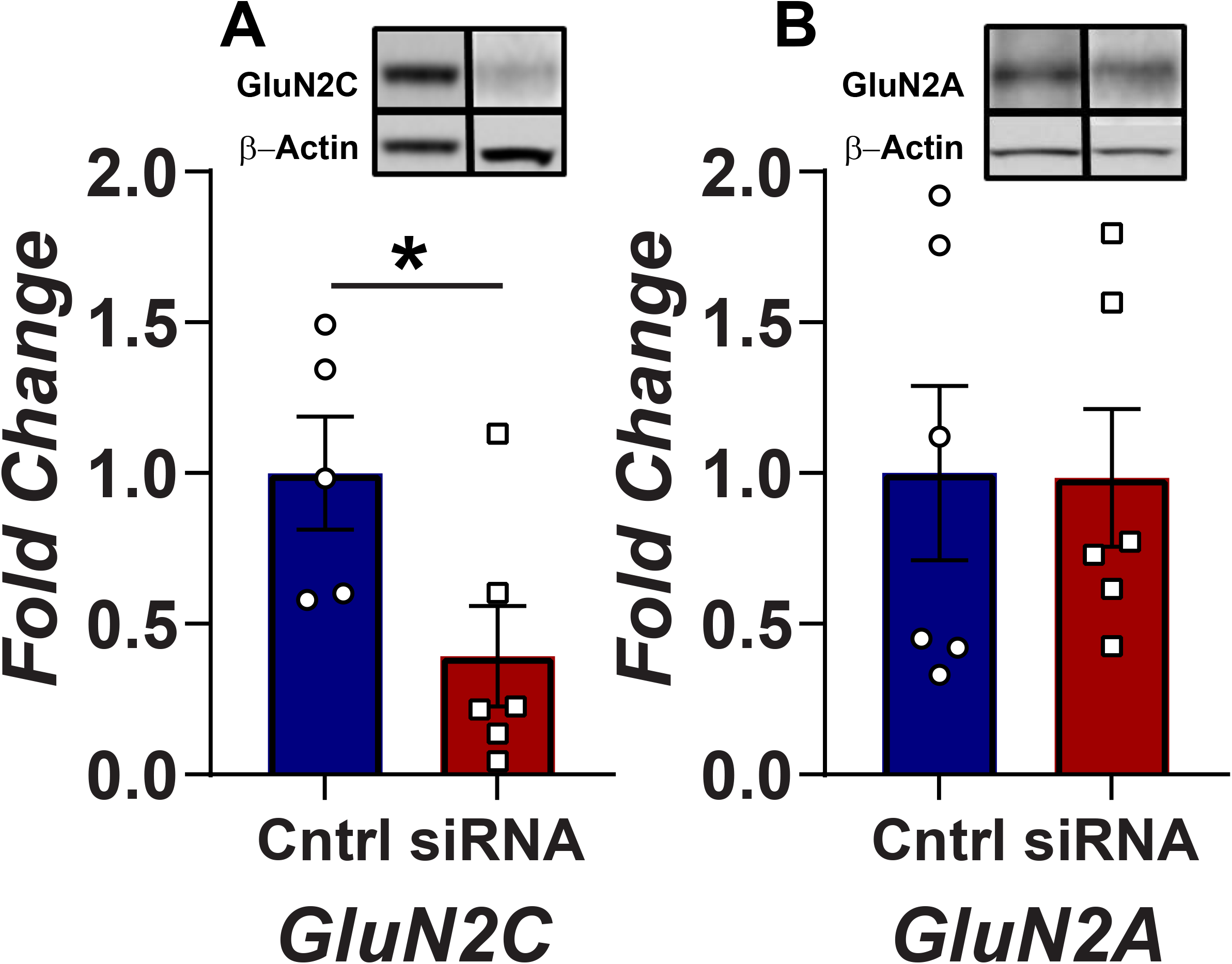
Grin2c intra-VTA siRNA injections significantly reduce GluN2C subunit protein levels in the VTA. **A)** siRNA microinjections against Grin2c subunit transcript produces a significant reduction in this subunit immunoreactivity. The histogram bars represent means +/- SEM of normalized NMDAR GluN2C subunit protein levels in animals injected with the Grin2c siRNA or the negative control siRNA (N=12 per condition) (*p <0.05; Student’s t-test) in VTA homogenates. The inset shows representative examples of signals from GluN2C and β-actin after Western blotting procedure. **B)** The histogram bars represent means +/- SEM of normalized NMDAR GluN2A subunit protein levels in animals injected with the Grin2c siRNA or the negative control siRNA (N=12 per condition) in VTA homogenates. The inset shows representative examples of signals from GluN2A and β-actin after Western blotting procedure.

### Down-regulation of the GluN2C subunit in VTA strongly reduces the Dorsal Raphe reward signal

The selective downregulation of the NMDAR GluN2C subunit produced a decrease in reward-seeking behavior. Figure 4 shows the behavioral data collected 24 h after the last siRNA microinjection for representative subjects. Intra-VTA microinjections of the active siRNA against Grin2c produced a rightward displacement of the curve that relates the nose-poke rate to the pulse frequency (R/F curve; Fig. 4C), reflecting a reduction in transmission of the reward signal within the VTA. On the contrary, VTA microinjections of the inactive RNA sequence did not affect the transmission of the reward signal as they did not produce a significant displacement of the R/F curve (Fig. 4A).

**Figure 4.**
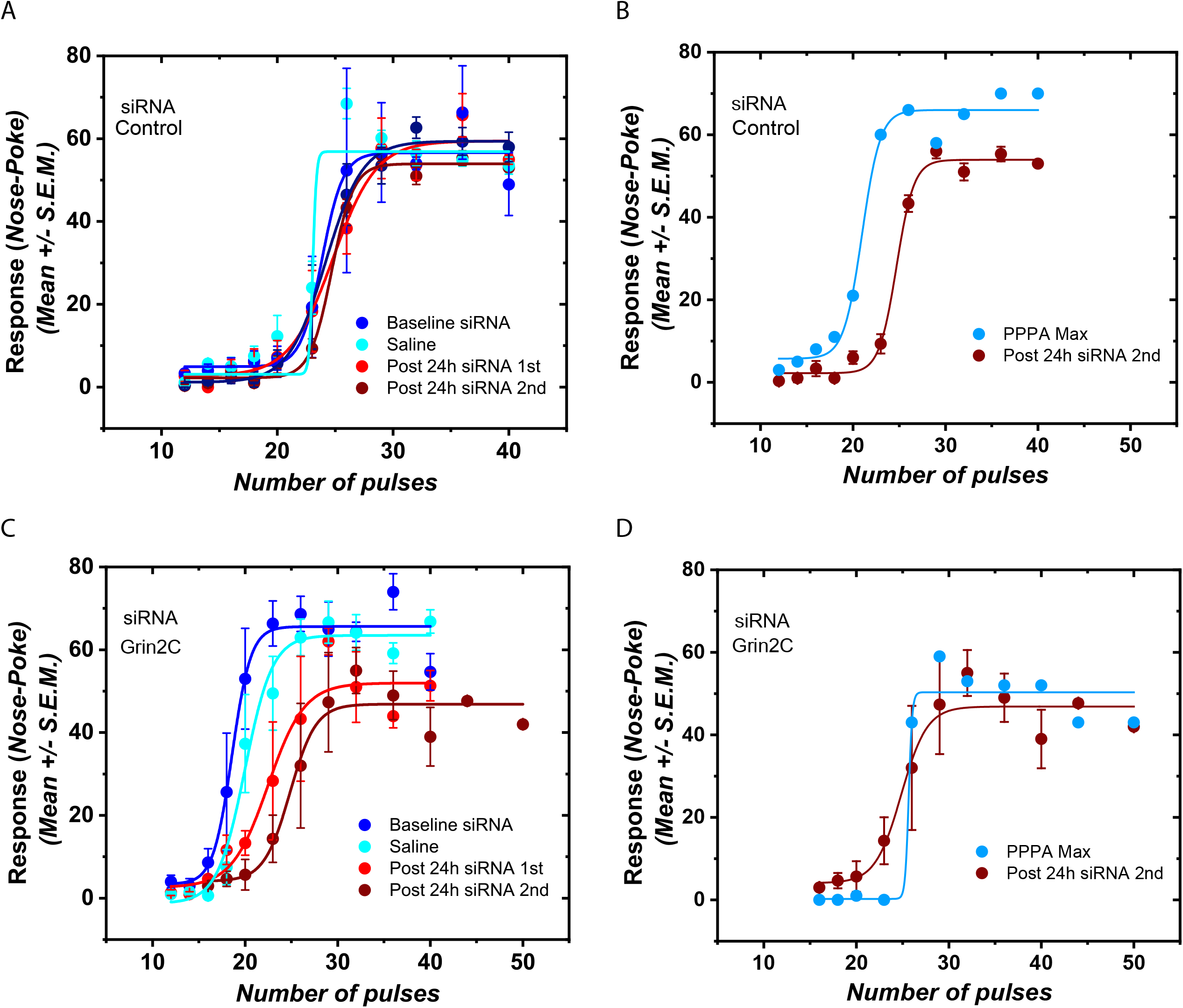
Intra-VTA Grin2c siRNA injection reduced rate-frequency response from dorsal raphe stimulation. The figure shows representative rate-frequency curves for selected subjects showing the effect of the different siRNA treatment and intra-VTA injection of PPPA (0.825 nmol/0.5μl/side). **A)** Treatment with the negative control siRNA (0.5 μg/side) produced no effect on the curve that relates the nose-poke rate with the electrical pulse trains. The baseline curve and the one obtained 24-h after the first and second siRNA injections overlap. **B)** In the animals treated with the negative control siRNA Intra-VTA injection with PPPA produced a leftward and upward shift of rate-frequency curve. **C)** Treatment with siRNA (0.5μg /side) against Grin2c produced a decrease of the rewarding effects that arise from dorsal raphe electrical stimulation. When contrasted against the baseline curves, the rate frequency curve obtained 24-h after the last siRNA injection is displaced to the right. A decrease in the upper asymptote is also observed. Interestingly, the Grin2c siRNA treatment alters PPPA effect. **D)** siRNA injections against Grin2c obliterate the effects of PPPA on brain stimulation reward and motor performance. The pretreatment with the Grin2c siRNA renders PPPA ineffective for modifying the reward signal that arises from dorsal raphe stimulation.

To further characterize the specific role of the GluN2C subunit in relaying the reward signal in VTA neurons, we used a pharmacological approach to the target NMDARs in animals that had received either a scrambled or Grin2c siRNA injection. First, in animals microinjected with the inactive siRNA, the NMDA antagonist, PPPA, enhanced the reward signal in the VTA; it produced a leftward displacement of the R/F curve (Fig. 4B) so that lower frequencies delivery to the DR were required to generate a reward. Interestingly, PPPA failed to alter the reward signal (no displacement of the R/F curve) in the animals microinjected with the active Grin2c siRNA (Fig. 4D).

Figure 5 shows the average changes in the M50 reward index and the maximum response. Intra-VTA injection of saline does not produce a significant change in the stimulation threshold (M50) [t_(19)_ = 1.696; p=0.10] or the maximum response rate [t_(21)_ = 1.009; p=0.3246] (Fig. 5 A-B). On the other hand, intra-VTA injection of siRNA against Grin2c produced a 38.17% increase (SEM = 7.12) in M50 when compared against their baseline, while the M50 of the rats that received the nonactive siRNA only changed by 1.5 % (SEM = 2.95). The increase in M50 observed in the subjects that received the siRNA against Grin2c is significantly different from that of the ones that received the scramble sequence (Fig. 5C) [t_(20)_ =4.754 p=0.0001] and reflects a significant reduction in transmission of DR reward signal to VTA neurons. Active siRNA also produced a small but significant 17.3% decrease in the maximum response rate (Fig. 5D) [t_(21)_ = 2.095; p=0.048].

**Figure 5.**
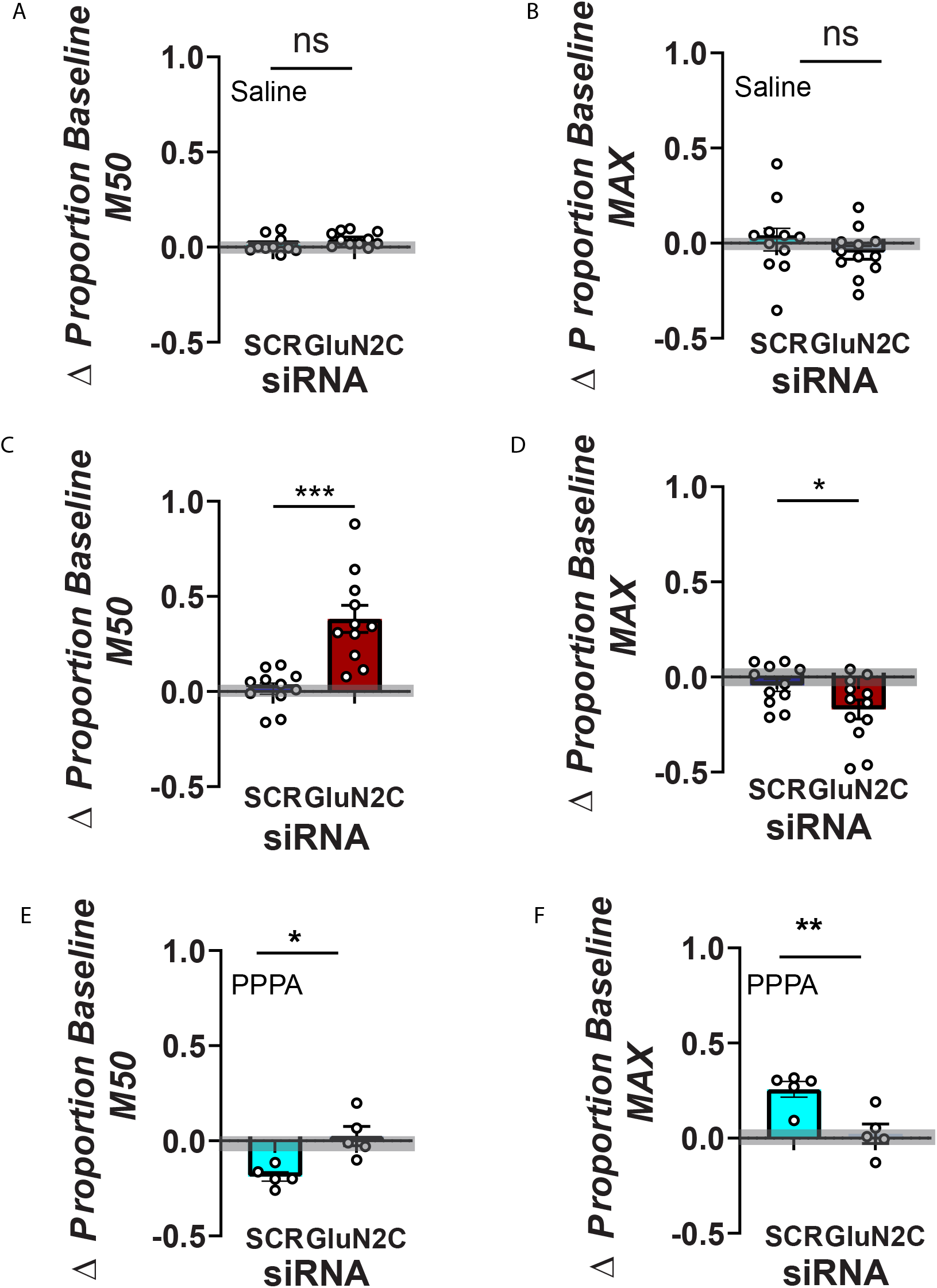
Shifts caused by control and Grin2c siRNAs in stimulation threshold and maximum response rate. **A-B)** Histogram bars represent means +/- SEM; saline injection in the VTA does not produces a reliable change in M50 nor MAX **C)** Treatment with the control siRNA did not produce a significant change in the stimulation threshold (M50), but the treatment with the siRNA against the Grin2c subunit produces a significant increase. **D)** The siRNA against Grin2c produced a significant decrease in the maximum response rate (MAX) compared to subjects that received the scramble siRNA. **E)** siRNA downregulation of the Grin2c subunit in the VTA abolishes the enhancing effect of PPPA. PPPA produced a decrease in the self-stimulation threshold only in those animals that were pre-treated with the scrambled siRNA. The self-stimulation threshold showed no reliable change in those animals pre-treated with the siRNA against Grin2c. The PPPA enhancement effect on M50 values in the scrambled siRNA group is significantly different from that of the group that received the siRNA against Grin2c. **F)** Similarly, PPPA only produced an increase in the maximum response rate in those animals that were pre-treated with the scrambled siRNA and not with the siRNA against Grin2C. The gray area denotes the 95% confidence interval of the baseline curves. (*p <0.05, ** p<0.01, *** p<0.0001; Student’s t-test).

The pharmacological manipulation of VTA glutamate using PPPA produced reliable changes in M50 and maximum response rate only in the animals that received the inactive siRNA treatment. PPPA enhanced the reward signal (reduction in M50 index, Fig. 5E). M50 decreased on average by 18.7 % (SEM = 2.38), and the maximum response rate increased on average by 25% (SEM = 4.15; Fig. 5F). For the subjects that received the siRNA against Grin2c, PPPA increased the M50 by 2% (SEM = 4.17) and the maximum response rate by 3.6% (SEM=4.4) (Fig. 5E and 5F). The differences between the inactive and active siRNA are significant for the changes produced in M50 for PPPA [t_(8)_ =3.782 p=0.005] as well as for the changes produced in the maximum rate PPPA [t_(8)_ =3.534; p=0.007]. Together, these data suggest that the GluN2C subunit is vital to relay the reward signal in VTA neurons.

## Discussion

Glutamate-containing terminals establish synaptic contacts with VTA DA neurons [36,37] and rewarding electrical stimulation is associated with an increase in VTA glutamate release [38]. Some of those VTA terminals come from the DR, which sends the highest number and density of glutamatergic projections [6].

Several studies have shown that NMDA receptors play a key role in activating DA neurons. Iontophoretic administration of NMDA receptor antagonists [39], as well as genetic deletion of the NMDA receptor GluN1 subunit from DA neurons, selectively attenuate DA phasic firing and reduces learning in cue-dependent learning tasks [40]. VTA downregulation of the GluN1 subunit also decreases brain stimulation reward [30]. Other research groups looking at depressive symptoms in rodents have shown that inactivation of the GluN1 subunit in VTA altered VTA dopamine cells, which correlated with behavioral deficits associated with anhedonia[41]. The fact that deletion or down-regulation of the GluN1 subunit produces profound effects is not surprising, as this subunit is a necessary component in forming functional NMDA receptor tetrameric complexes [42,43].

Here, we first measured Grin2 subunit transcript levels among several limbic brain regions involved in motivated behaviors. The relative magnitude of each Grin2 mRNA level varied among regions. Grin2a and Grin2b are the most abundant in the forebrain and diencephalic regions, while Grin2d is the least abundant, findings that are consistent with previous reports (Ishii et al., 1993; Monyer et al 1994). In more caudal mesencephalic regions such as VTA and DR, we found the Grin2c is the most abundant, and the levels of Grin2d is much higher than in forebrain regions and the hippocampus. These findings are also consistent we previous results (Fasulo and Hemby, 2003; Ishii et al 1993; Monyer et al., 1994). These results suggest an important and underappreciated role for the NMDAR containing the GluN2C subunit in the limbic functions of these mesencephalic brain areas.

Another major finding of our study is that the NMDA receptor containing the GluN2C subunit within the VTA is a necessary component for the transmission of the rewarding signal arising from DR. Because the cellular machinery required for silencing the targeted RNA is primarily located in the nucleus of a cell, VTA infusion of siRNA is expected to initially downregulate receptors expressed on cell bodies located within the VTA, interfering with the mRNA and the processes required for protein translation. Within the first 24h after the infusion, it must be less effective at reducing receptors located on afferent terminals. Our selective GluN2C subunit downregulation *via* siRNA produced a significant reduction in brain stimulation reward and a change in the performance capacity; effects were detected 24h after the first infusion. These reductions are inferred by the rightward and downward (reduction in asymptote) displacement of the R/F curve [44]. Both effects could be associated with a reduction in the release of DA from the nucleus accumbens since this decrease can affect both reward and motor function [45]. It has been shown that electrical and optical stimulation of DR neurons produces an increase in DA cell firing and DA phasic release [2,15,46] via glutamatergic DR efferents to the VTA [1,2,20]. Substantial reduction in the GluN2C input likely decreases glutamatergic excitability in DA neurons, reducing the overall DA release in terminal areas. The reduction of brain stimulation reward is specific to GluN2C since a previous study showed that downregulation of GluN2A or/and GluN2D did not produce a significant change in the curve that relates stimulation of the DR with nose-poke behavior [30].

Our double fluorescent hybridization data show that a substantial amount of Grin2c mRNA is colocalized in VTA TH+ cells. This observation is in line with single-cell expression profiling on DA neurons, which shows that the Grin2c mRNA is expressed in mesocorticolimbic DA neurons [47–49]. Using qRT-PCR, we observed that Grin2c mRNA is the most abundantly expressed in DR and VTA compared to other Grin2 subunit transcripts. In fact, Grin2c mRNA is expressed in all areas of the brain observed in the adult rat brain, but its relative expression changes accordingly to where it is measured in the anterior-posterior axis. In comparison, the Grin2a and Grin2b transcripts are the most abundant mRNAs in the anterior and medial parts of the brain, the balance shifts towards Grin2c in posterior brain areas. This observation is significant because, until recently, not many studies have been done regarding the role of this NMDA receptor subunit, which was believed to be confined to the cerebellum, thalamus, and olfactory bulb and to be present at very low levels in the hippocampus of the adult rodent brain [50,51]. However, present results and recent observations using novel reporter mice [52] suggest this is not the case.

Moreover, there is accumulating evidence that the NMDA receptor containing the GluN2C subunit plays an important role in several domains. For example, NMDA receptors containing GluN2C are required for the acquisition of conditioned fear and working memory because Grin2c knockout mice show deficits in associating cues with aversive consequences, as well as navigating an eight-arm radial maze [53]. Also, they spend more time immobile in the forced swim test [54]. At the cortical level, this receptor controls the balance of excitatory and inhibitory activity in the medial prefrontal cortex since Grin2c knockout mice have reduced excitatory postsynaptic currents, increased inhibitory postsynaptic currents, and reduced spine density [55].

Can a downregulation of Grin2c mRNA in VTA TH-cells contribute to the reduction in reward observed in this study? It is well known that VTA interneurons provide a strong, tonic, inhibitory drive onto DA neurons. A reduction in glutamate excitatory input to these inhibitory neurons allows DA cells to go from a silent state into a tonic firing state, increasing the likelihood that they will fire in phasic mode. Such a change in DA firing mode is expected to translate into an enhancement of reward rather than the attenuation we observed. Therefore, it is unlikely that GluN2C-containing NMDA receptors containing GluN2C in these TH-cells play a role in relaying the reward signal.

To further characterize the effect of down-regulation of the GluN2C receptor in TH+ cells, we injected PPPA, an NMDA receptor antagonist, into the VTA. Previous studies from our laboratory [5,30,32] have shown that PPPA enhances brain stimulation reward, an effect that could be explained by a reduction in the excitatory drive to VTA inhibitory interneurons. Consistent with our previous results, PPPA produced a reliable enhancement in reward and performance; however, these effects were only observed in the animals that received the inactive siRNA; in those animals where the GluN2C subunit was downregulated, PPPA produced no change in brain stimulation reward. These results suggest that Glun2C subunit are also involved in the excitatory drive to VTA inhibitory interneurons; the reduction in the reward that we observed following local infusion of Grin2c siRNA must result from the net effect of disinhibition and excitation of VTA DA signaling; therefore, unable to substantially change brain stimulation reward.

### Technical Considerations

It has been demonstrated that the detection of specific reward changes in the presence of a decrease in operant response cannot be achieved by the unique measure of the response rate. To circumvent this problem, we applied the psychophysical technique (curve shift method), a technique that allows us to determine a reliable index of brain stimulation reward (M50) when used with the appropriate parameters (0.1 ms cathodal pulses, fixed train duration, and fixed current intensity) of electrical stimulation (Gallistel et al., 1980). The M50 index is indeed sensitive to changes in rates of responding, but as shown by Miliaressis et al (1985) and Fouriezos et al (1990), the mean 50% increase in M50 that we measured after injection of the active siRNA cannot be due to the mean 15% decrease in maximum rates of responding that we also observed; it can only be the consequence of a specific reward attenuation. Although electrical stimulation does not allow activating a selective population of neurons, it is used here with a psychophysical method that permits us to detect attenuation of brain stimulation reward that results from a reduction in VTA GluN2C subunits.

## Conclusion

We demonstrated that a localized decrease in GluN2C-containing NMDA receptors in the VTA significantly alters the rewarding signal originated in the DR. This observation adds to other studies suggesting that the GluN2C receptor plays an important role in several disorders, including drug abuse [56] and schizophrenia [53,55]. Furthermore, our results suggest that the levels of this receptor in the VTA might be at the root of anhedonia.

## Acknowledgments

The present research was supported by NSERC grant # 119057 to Pierre-Paul Rompré, Herbert H. Jasper fellowship to Giovanni Hernandez, CIHR operating grant #130407 to Daniel Lévesque.

## Figure Captions

**Supplementary Figure 1: Schematic of the experimental behavioral procedure**.

**Supplementary Figure 2: Low-level magnification of Double FISH immunodetection**.

**Supplementary Table 1: Primer sequences for qRT-PCR Grin2a-d analysis**

## References

[1] Mcdevitt RA, Tiran-cappello A, Harvey BK, Bonci A, Shen H, Balderas I, et al. Serotonergic versus Nonserotonergic Dorsal Raphe Projection Neurons1: Differential Participation in Reward Article Serotonergic versus Nonserotonergic Dorsal Raphe Projection Neurons1: Differential Participation in Reward Circuitry 2014:1857–69.

[2] Qi J, Zhang S, Wang H-L, Wang H, de Jesus Aceves Buendia J, Hoffman AF, et al. A glutamatergic reward input from the dorsal raphe to ventral tegmental area dopamine neurons. Nat Commun 2014;5:5390. https://doi.org/10.1038/ncomms6390.

[3] Boye SM, Rompré P-P. Mesencephalic substrate of reward: Axonal connections. J Neurosci 1996;16:3511–20.

[4] Hernandez G, Khodami-Pour A, Lévesque D, Rompré PP. Reduction in ventral midbrain NMDA receptors reveals two opposite modulatory roles for glutamate on reward. Neuropsychopharmacology 2015;40:1682–91. https://doi.org/10.1038/npp.2015.14.

[5] Bergeron S, Rompré P-P. Blockade of ventral midbrain NMDA receptors enhances brain stimulation reward: a preferential role for GluN2A subunits. Eur Neuropsychopharmacol 2013;23:1623–35. https://doi.org/10.1016/j.euroneuro.2012.12.005.

[6] Watabe-Uchida M, Zhu L, Ogawa SK, Vamanrao A, Uchida N. Whole-brain mapping of direct inputs to midbrain dopamine neurons. Neuron 2012;74:858–73. https://doi.org/10.1016/j.neuron.2012.03.017.

[7] Wise R a. Dopamine, learning and motivation. Nat Rev Neurosci 2004;5:483–94. https://doi.org/10.1038/nrn1406.

[8] Hernandez G, Breton Y-A, Conover K, Shizgal P. At what stage of neural processing does cocaine act to boost pursuit of rewards? PLoS One 2010;5:e15081. https://doi.org/10.1371/journal.pone.0015081.

[9] Hernandez G, Trujillo-Pisanty I, Cossette M-P, Conover K, Shizgal P. Role of Dopamine Tone in the Pursuit of Brain Stimulation Reward. J Neurosci 2012;32:11032–41. https://doi.org/10.1523/JNEUROSCI.1051-12.2012.

[10] Schultz W. A Neural Substrate of Prediction and Reward. Science (80-) 1997;275:1593–9. https://doi.org/10.1126/science.275.5306.1593.

[11] Salamone JD. Behavioral pharmacology of dopamine systems: A new synthesis. mesolimbic dopamine Syst. From Motiv. to action, 1991, p. 599–614.

[12] Wise R a, Rompre PP. Brain dopamine and reward. Annu Rev Psychol 1989;40:191–225. https://doi.org/10.1146/annurev.ps.40.020189.001203.

[13] Simon H, Le Moal M, Cardo B. Intracranial self-stimulation from the dorsal raphe nucleus of the rat: effects of the injection of para-chlorophenylalanine and of alpha-methylparatyrosine. Behav Biol 1976. https://doi.org/10.1016/S0091-6773(76)91486-3.

[14] Rompre PP, Miliaressis E. Pontine and mesencephalic substrates of self-stimulation. Brain Res 1985;359:246–59.

[15] Moisan J, Rompre PP. Electrophysiological evidence that a subset of midbrain dopamine neurons integrate the reward signal induced by electrical stimulation of the posterior mesencephalon. Brain Res 1998;786:143–52. https://doi.org/doi.10.1016/S0006-8993(97)01457-1.

[16] Jacobs BL, Azmitia EC. Structure and function of the brain serotonin system. Physiol Rev 1992;72:165–230. https://doi.org/10.1152/physrev.1992.72.1.165.

[17] Vasudeva RK, Lin RCS, Simpson KL, Waterhouse BD. Functional organization of the dorsal raphe efferent system with special consideration of nitrergic cell groups. J Chem Neuroanat 2011;41:281–93. https://doi.org/10.1016/j.jchemneu.2011.05.008.

[18] Wang RY, Aghajani GK. Antidromically identified serotonergic neurons in the rat midbrain raphe: evidence for collateral inhibition. Brain Res 1977. https://doi.org/10.1016/0006-8993(77)90719-3.

[19] Rompré PP, Miliaressis E. Behavioral determination of refractory periods of the brainstem substrates of self-stimulation. Behav Brain Res 1987;23:205–19.

[20] Liu Z, Zhou J, Li Y, Hu F, Lu Y, Ma M, et al. Dorsal Raphe Neurons Signal Reward through 5-HT and Glutamate. Neuron 2014;81:1360–74. https://doi.org/10.1016/j.neuron.2014.02.010.

[21] Sesack SR, Carr DB, Omelchenko N, Pinto A. Anatomical Substrates for Glutamate-Dopamine Interactions. Ann N Y Acad Sci 2003;1003:36–52. https://doi.org/10.1196/annals.1300.066.

[22] Mogenson GJ, Jones DL, Yim CY. From motivation to action: functional interface between the limbic system and the motor system. Prog Neurobiol 1980;14:69–97. https://doi.org/10.1016/0301-0082(80)90018-0.

[23] Chergui K, Charléty PJ, Akaoka H, Saunier CF, Brunet J -L, Buda M, et al. Tonic Activation of NMDA Receptors Causes Spontaneous Burst Discharge of Rat Midbrain Dopamine Neurons In Vivo. Eur J Neurosci 1993;5:137–44. https://doi.org/10.1111/j.1460-9568.1993.tb00479.x.

[24] Grace AA, Bunney BS. The control of firing pattern in nigral dopamine neurons: burst firing. J Neurosci 1984;4:2877–90. https://doi.org/6150071.

[25] Hyland BI, Reynolds JNJ, Hay J, Perk CG, Miller R. Firing modes of midbrain dopamine cells in the freely moving rat. Neuroscience 2002;114:475–92. https://doi.org/10.1016/S0306-4522(02)00267-1.

[26] Suaud-Chagny MF, Chergui K, Chouvet G, Gonon F. Relationship between dopamine release in the rat nucleus accumbens and the discharge activity of dopaminergic neurons during local in vivo application of amino acids in the ventral tegmental area. Neuroscience 1992. https://doi.org/10.1016/0306-4522(92)90076-E.

[27] Tsai H-C, Zhang F, Adamantidis A, Stuber GD, Bonci A, de Lecea L, et al. Phasic firing in dopaminergic neurons is sufficient for behavioral conditioning. Science 2009;324:1080–4. https://doi.org/10.1126/science.1168878.

[28] Pérez-Otaño I, Schulteis CT, Contractor A, Lipton SA, Trimmer JS, Sucher NJ, et al. Assembly with the NR1 subunit is required for surface expression of NR3A-containing NMDA receptors. J Neurosci 2001;21:1228–37. https://doi.org/10.1523/jneurosci.21-04-01228.2001.

[29] McIlhinney RAJ, L. Bourdellès B, Molnár E, Tricaud N, Streit P, Whiting PJ. Assembly intracellular targeting and cell surface expression of the human N-methyl-D-aspartate receptor subunits NR1a and NR2A in transfected cells. Neuropharmacology 1998;37:1355–67. https://doi.org/10.1016/S0028-3908(98)00121-X.

[30] Hernandez G, Khodami-Pour A, Lévesque D, Rompré P-PP. Reduction in Ventral Midbrain NMDA Receptors Reveals Two Opposite Modulatory Roles for Glutamate on Reward. Neuropsychopharmacology 2015;40:1682–91. https://doi.org/10.1038/npp.2015.14.

[31] Salahpour A, Medvedev IO, Beaulieu J-M, Gainetdinov RR, Caron MG. Local knockdown of genes in the brain using small interfering RNA: a phenotypic comparison with knockout animals. Biol Psychiatry 2007;61:65–9. https://doi.org/10.1016/j.biopsych.2006.03.020.

[32] Ducrot C, Fortier E, Bouchard C, Rompré P-P. Opposite modulation of brain stimulation reward by NMDA and AMPA receptors in the ventral tegmental area. Front Syst Neurosci 2013;7:57. https://doi.org/10.3389/fnsys.2013.00057.

[33] Vandesompele J, De Preter K, Pattyn F, Poppe B, Van Roy N, De Paepe A, et al. Accurate normalization of real-time quantitative RT-PCR data by geometric averaging of multiple internal control genes. Genome Biol 2002;3:RESEARCH0034. https://doi.org/10.1186/gb-2002-3-7-research0034.

[34] Lein ES, Hawrylycz MJ, Ao N, Ayres M, Bensinger A, Bernard A, et al. Genome-wide atlas of gene expression in the adult mouse brain. Nature 2007;445:168–76. https://doi.org/10.1038/nature05453.

[35] Allen Institute for Brain Science. Allen Mouse Brain Atlas 2010. https://mouse.brain-map.org/search/index.

[36] Carr DB, Sesack SR. Projections from the rat prefrontal cortex to the ventral tegmental area: target specificity in the synaptic associations with mesoaccumbens and mesocortical neurons. J Neurosci 2000;20:3864–73.

[37] Omelchenko N, Sesack SR. Ultrastructural analysis of local collaterals of rat ventral tegmental area neurons: GABA phenotype and synapses onto dopamine and GABA cells. Synapse 2009;63:895–906. https://doi.org/10.1002/syn.20668.

[38] You ZB, Chen YQ, Wise RA. Dopamine and glutamate release in the nucleus accumbens and ventral tegmental area of rat following lateral hypothalamic self-stimulation. Neuroscience 2001;107:629–39. https://doi.org/10.1016/S0306-4522(01)00379-7.

[39] Overton P, Clark D. Iontophoretically administered drugs acting at the N-methyl-D-aspartate receptor modulate burst firing in A9 dopamine neurons in the rat. Synapse 1992;10:131–40. https://doi.org/10.1002/syn.890100208.

[40] Zweifel LS, Parker JG, Lobb CJ, Rainwater A, Wall VZ, Fadok JP, et al. Disruption of NMDAR-dependent burst firing by dopamine neurons provides selective assessment of phasic dopamine-dependent behavior. Proc Natl Acad Sci 2009;106:7281–8. https://doi.org/10.1073/pnas.0813415106.

[41] Jastrzebska K, Walczak M, Cieslak PE, Szumiec L, Turbasa M, Engblom D, et al. Loss of NMDA receptors in dopamine neurons leads to the development of affective disorder-like symptoms in mice. Sci Rep 2016;6:1–11. https://doi.org/10.1038/srep37171.

[42] Monyer H, Sprengel R, Schoepfer R, Herb A, Higuchi M, Lomeli H, et al. Heteromeric NMDA Receptors: Molecular and Functional Distinction of Subtypes. Science (80-) 1992;256:1217–21. https://doi.org/10.1126/science.256.5060.1217.

[43] Nakanishi S. Molecular diversity of glutamate receptors and implications for brain function. Science (80-) 1992;258:597–603. https://doi.org/10.1126/science.1329206.

[44] Miliaressis E, Rompre PP, Laviolette P, Philippe L, Coulombe D. The curve-shift paradigm in self-stimulation. Physiol Behav 1986;37:85–91. https://doi.org/10.1016/0031-9384(86)90388-4.

[45] Stellar JR, Kelley a E, Corbett D. Effects of peripheral and central dopamine blockade on lateral hypothalamic self-stimulation: evidence for both reward and motor deficits. Pharmacol Biochem Behav 1983;18:433–42.

[46] Hernandez G, Cossette MP, Shizgal P, Rompré PP. Ventral midbrain NMDA receptor blockade: From enhanced reward and dopamine inactivation. Front Behav Neurosci 2016;10:1–11. https://doi.org/10.3389/fnbeh.2016.00161.

[47] Anderegg A, Poulin JF, Awatramani R. Molecular heterogeneity of midbrain dopaminergic neurons - Moving toward single cell resolution. FEBS Lett 2015;589:3714–26. https://doi.org/10.1016/j.febslet.2015.10.022.

[48] Saunders A, Macosko EZ, Wysoker A, Goldman M, Krienen FM, de Rivera H, et al. Molecular Diversity and Specializations among the Cells of the Adult Mouse Brain. Cell 2018;174:1015-1030.e16. https://doi.org/10.1016/j.cell.2018.07.028.

[49] Poulin JF, Gaertner Z, Moreno-Ramos OA, Awatramani R. Classification of Midbrain Dopamine Neurons Using Single-Cell Gene Expression Profiling Approaches. Trends Neurosci 2020;43:155– 69. https://doi.org/10.1016/j.tins.2020.01.004.

[50] Monyer H, Burnashev N, Laurie DJ, Sakmann B, Seeburg PH. Developmental and regional expression in the rat brain and functional properties of four NMDA receptors. Neuron 1994;12:529–40. https://doi.org/10.1016/0896-6273(94)90210-0.

[51] Wenzel A, Scheurer L, Künzi R, Fritschy JM, Mohler H, Benke D. Distribution of NMDA receptor subunit proteins NR2A, 2B, 2C and 2D in rat brain. Neuroreport 1995;7:45–8.

[52] Ravikrishnan A, Gandhi PJ, Shelkar GP, Liu J, Pavuluri R, Dravid SM. Region-specific Expression of NMDA Receptor GluN2C Subunit in Parvalbumin-Positive Neurons and Astrocytes: Analysis of GluN2C Expression using a Novel Reporter Model. Neuroscience 2018;380:49–62. https://doi.org/10.1016/j.neuroscience.2018.03.011.

[53] Hillman BG, Gupta SC, Stairs DJ, Buonanno A, Dravid SM. Behavioral analysis of NR2C knockout mouse reveals deficit in acquisition of conditioned fear and working memory. Neurobiol Learn Mem 2011;95:404–14. https://doi.org/10.1016/j.nlm.2011.01.008.

[54] Shelkar GP, Pavuluri R, Gandhi PJ, Ravikrishnan A, Gawande DY, Liu J, et al. Differential effect of NMDA receptor GluN2C and GluN2D subunit ablation on behavior and channel blocker-induced schizophrenia phenotypes. Sci Rep 2019;9:1–11. https://doi.org/10.1038/s41598-019-43957-2.

[55] Gupta SC, Ravikrishnan A, Liu J, Mao Z, Pavuluri R, Hillman BG, et al. The NMDA receptor GluN2C subunit controls cortical excitatory-inhibitory balance, neuronal oscillations and cognitive function. Sci Rep 2016;6:1–13. https://doi.org/10.1038/srep38321.

[56] Joffe ME, Grueter BA. Cocaine Experience Enhances Thalamo-Accumbens N-Methyl-D-Aspartate Receptor Function. Biol Psychiatry 2016;80:671–81. https://doi.org/10.1016/j.biopsych.2016.04.002.

