## Supplementary figures and images for "Dorsal raphe stimulation relays a reward signal to the ventral tegmental area via GluN2C NMDA receptors"

### Supplementary fig 1

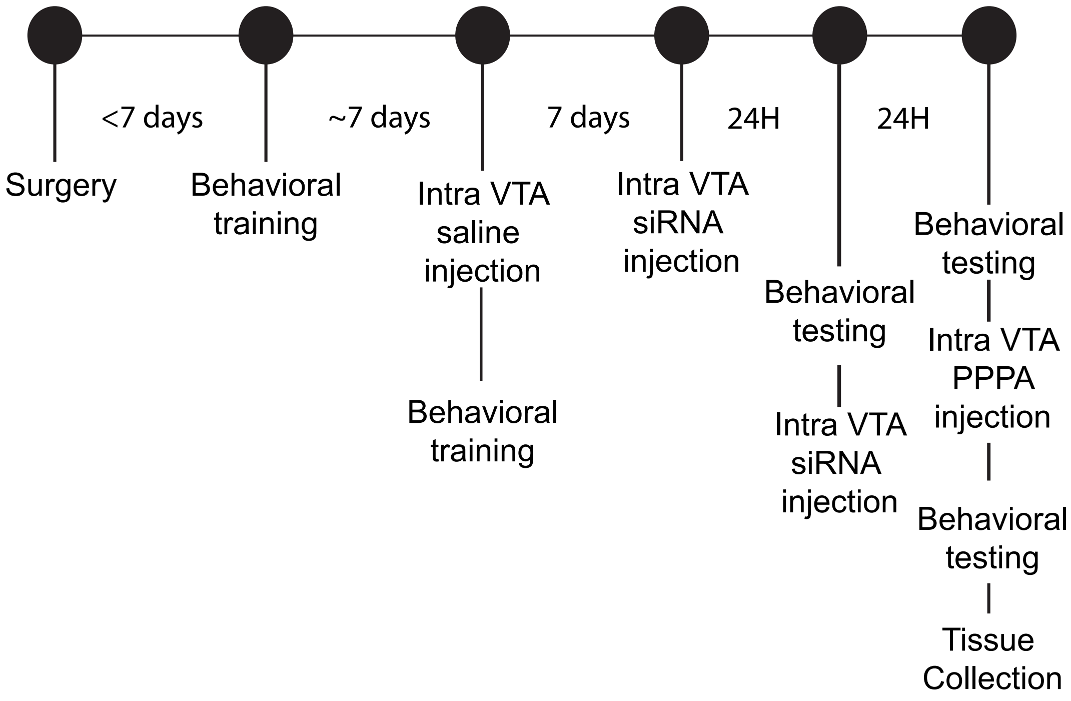

### Supplementary fig 2

TH

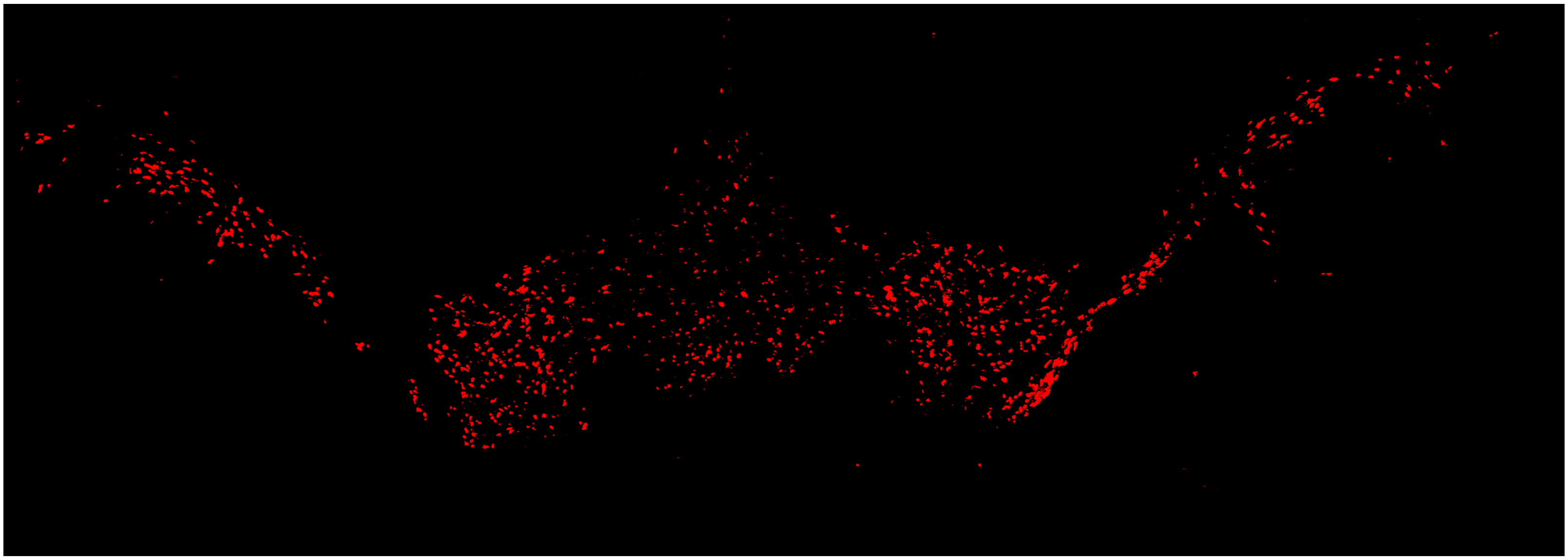

GluN2C

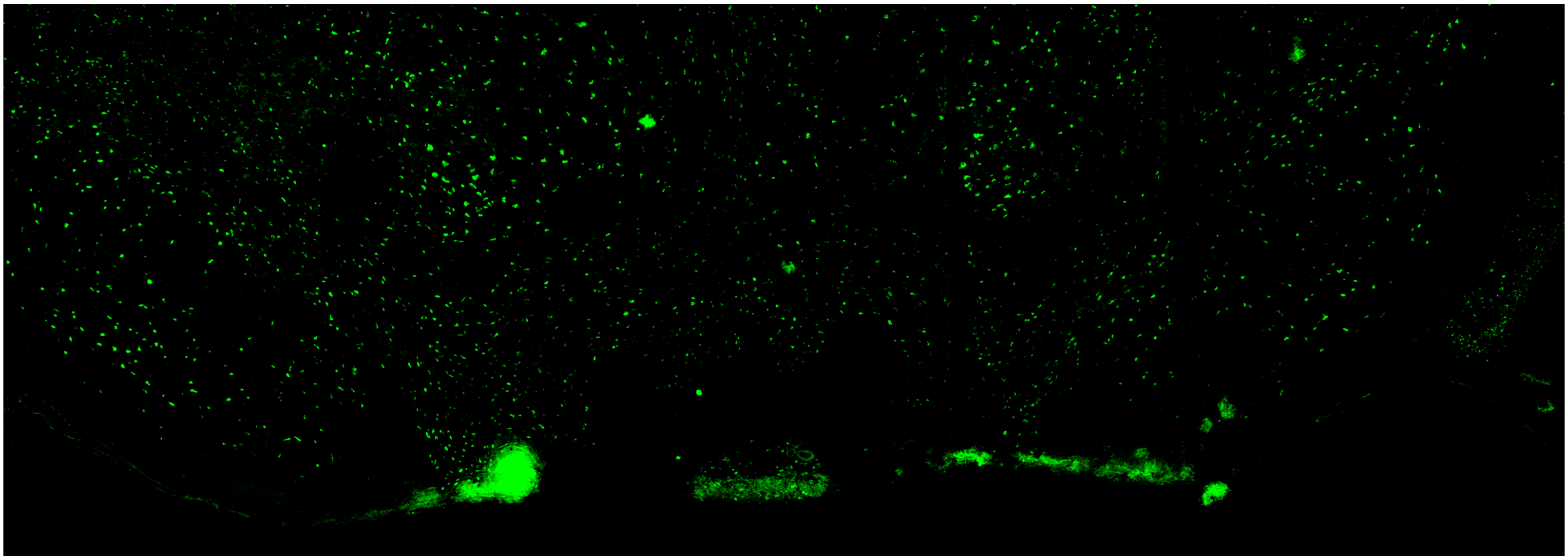

Merge

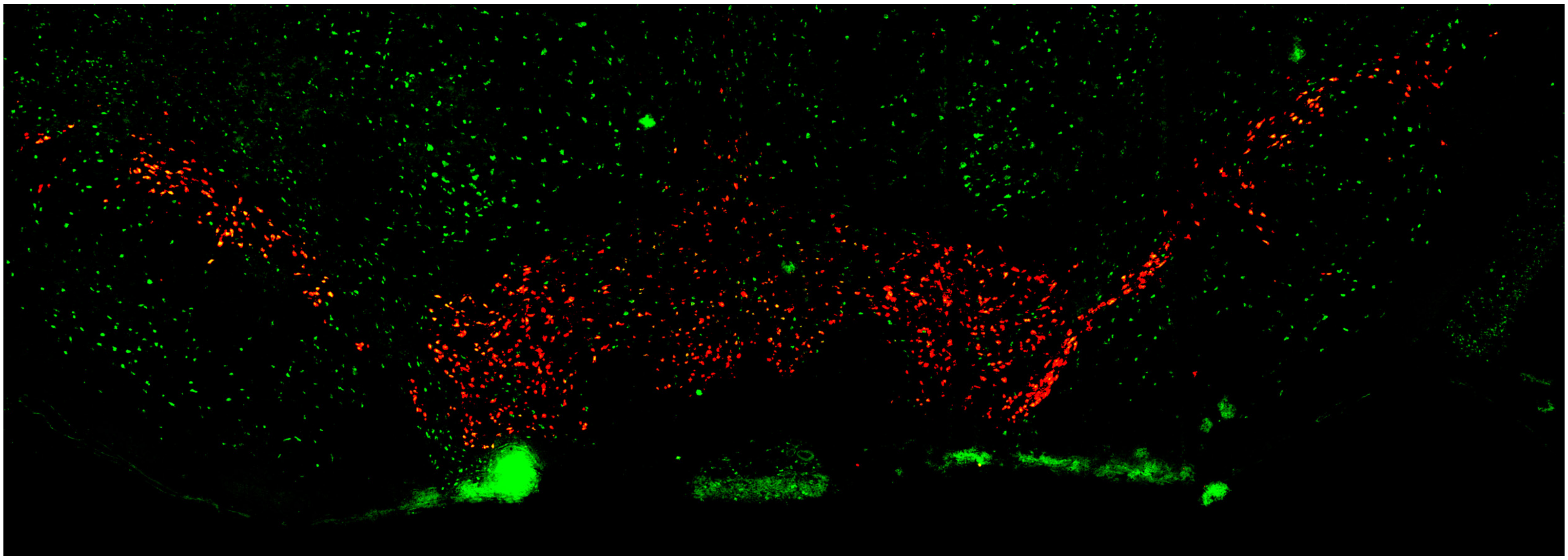
