## Supplementary table 1 for "Dorsal raphe stimulation relays a reward signal to the ventral tegmental area via GluN2C NMDA receptors"

**Primer sequences for qPCR Grin2 analysis**

|  | **Gene Symbol** |  | **Fwd** | **Rev** | **RefSeq** | | **Efficiency %** | |
| --- | --- | --- | --- | --- | --- | --- | --- | --- |
|  | Grin2a |  | atgatcatggctgacaagga | cagcataactgtggcttgct | | NM_012573.3 | | 102 |
|  | Grin2b |  | catggtatctcgcagcaatg | cagcgctgaatggctctaa | | NM_012574.1 | | 91 |
|  | Grin2c |  | ggcactcctgcaacttctg | gttctggcagatccctgaga | | NM_012575.3 | | 100 |
|  | Grin2d |  | gcccattctcgacttcctg | aggaaggtggagcccttct | | NM_022797.1 | | 107 |
|  | Gapdh |  | ccctcaagattgtcagcaatg | agttgtcatggatgaccttgg | | NM_017008.3 | | 97 |
|  | Hprt |  | gaccggttctgtcatgtcg | acctggttcatcatcactaatcac | | NM_012583 | | 105 |
